# Alternative splicing of *SORBS1* affects neuromuscular junction formation and stability in myotonic dystrophy type 1

**DOI:** 10.1101/2024.12.23.630072

**Authors:** Caroline Hermitte, Hortense de Calbiac, Gilles Moulay, Antoine Mérien, Jeanne Lainé, Hélène Polvèche, Michel Cailleret, Stéphane Vassilopoulos, Edor Kabashi, Denis Furling, Cécile Martinat, Morgan Gazzola

## Abstract

Myotonic dystrophy type 1 (DM1) is a multisystemic neuromuscular disease characterized by a CTG repeat expansion in the 3’ untranslated region of the gene coding for the *dystrophia myotonica protein kinase* (DMPK). Presence of expanded CTG repeats in DMPK-mRNAs leads to the sequestration of RNA binding factors such as the Muscleblind like (MBNL) proteins resulting in widespread splicing defects contributing to progressive muscle weakness and myotonia. Here, we show that abnormal splicing of *SORBS1* exon 25 found in skeletal muscle of myotonic dystrophy type 1 patients is a critical contributor to neuromuscular junction (NMJ) formation and maintenance. Forced exclusion of *SORBS1* exon 25 in mice results in NMJ degeneration with marked denervation and postsynaptic destabilization. In zebrafish, misregulation of *sorbs1* exon 25 results in reduced motor function and abnormal AChR cluster morphology. Finally, we observed that forcing *SORBS1* exon 25 exclusion in hiPSC-derived skeletal muscle cells reduces the formation of large AChR clusters upon agrin stimulation. Thus, our study identifies MBNL regulated *SORBS1* alternative splicing during skeletal muscle development as a critical event for NMJ formation and maintenance. The aberrant splicing of *SORBS1* exon 25 in DM1 expands our understanding of how splicing dysregulation compromises neuromuscular system communication, shedding light on the broader impact of mRNA splicing regulation on NMJ integrity.

## INTRODUCTION

Myotonic dystrophy type 1 (DM1), with a prevalence of 1 over 8000 individuals, is considered as one of the most common neuromuscular disorder worldwide^1^. This pathology is characterized by muscular alterations such as myotonia, muscle atrophy and muscle weakness^2^. DM1 is caused by the presence of CTG repeat expansions in the 3’ untranslated region (UTR) of the gene coding for the *dystrophia myotonica protein kinase* (DMPK). The severity of this disease correlates with the number of CTG repeats, ranging from congenital forms (>1500 repetitions) to adult forms and late onset (50-150 repetitions). Once transcribed, the CUG repeats form hairpin structures that sequester RNA binding factors such as Muscleblind like (MBNL) proteins, resulting in the formation of toxic foci within the cell nuclei^3,4^.

The sequestration of MBNL proteins results in widespread misregulation of alternative splicing including cassette exon, mutually exclusive exon, and retained introns^5^. To date, only a few mis-splicing events affecting genes such as *CLCN1*, *CLTC*, *BIN1* and *DMD* have been characterized at the pathological level and shown to contribute to muscle weakness and myotonia^6–11^. Additional splicing misregulation events have been identified in DM1, but their impact on muscle function remains largely unexplored^12^. Among them, mis-regulation of SORBS1 exon 25 splicing has also been reported in DM1 mouse models, including DMSXL and MBNL1 deficient mice^13–16^. We recently identified *SORBS1* exon 25 as one of the most significantly differentially spliced exons (DSEs) in DM1-affected human induced pluripotent stem cells (hiPSC) derived skeletal muscle cells, and in cells lacking MBNL isoforms 1 and 2 of MBNL (MBNL DKO)^17^.

The *SORBS1* gene encodes a protein involved in cytoskeleton modulation through interactions with paxillin, talin, and vinculin^18–20^, and has recently been described as implicated in acetylcholine receptor (AChR) cluster formation in mouse skeletal muscle cells^21^. These observations suggest that *SORBS1* splicing mis-regulation may play a previously unrecognized role in NMJ formation and stability. Given the critical role of NMJs in muscle contractions and locomotion, we hypothesized that mis-splicing of *SORBS1* exon 25 could significantly impair NMJ formation and maintenance in DM1. To explore this, the impact of *SORBS1* exon 25 splicing misregulation on NMJ integrity was investigated.

By utilizing forced exon-skipping strategies across diverse *in vitro* and *in vivo* models, including hiPSC-derived skeletal muscle cells, mice and zebrafish, our findings demonstrate that the SORBS1 protein encoded from the transcript containing exon 25 is crucial for NMJ formation and stability. Furthermore, the mis-regulation of SORBS1 exon 25 alternative splicing may play a role in pathological motor impairments.

## RESULTS

### *SORBS1* exon 25 is mis-regulated in skeletal muscle from DM1 patients

In our previous study, we identified widespread alternative splicing misregulation in both DM1 and MBNL1/2 double knockout (DKO) hiPSC-derived skeletal muscle cells^17^. This analysis revealed 254 DSEs across control, DM1, and DKO conditions. Notably, *SORBS1* exon 25 exhibited the most significant downregulation among these DSEs, with a marked reduction in the percentage of splice inclusion (ΔPSI) of 68% in DM1 myotubes compared to controls (**Fig. 1a**). RTqPCR analyses performed in control, DM1, and DKO hiPSC-derived skeletal muscle cells corroborated this finding, with a ΔPSI of 81 ± 1.8% in DM1 skeletal muscle cells and an even more pronounced reduction of 86.5 ± 0.7% in DKO skeletal muscle cells, confirming the regulatory influence of MBNL1/2 proteins on *SORBS1* splicing (**Fig. 1b**).

**Figure 1.**
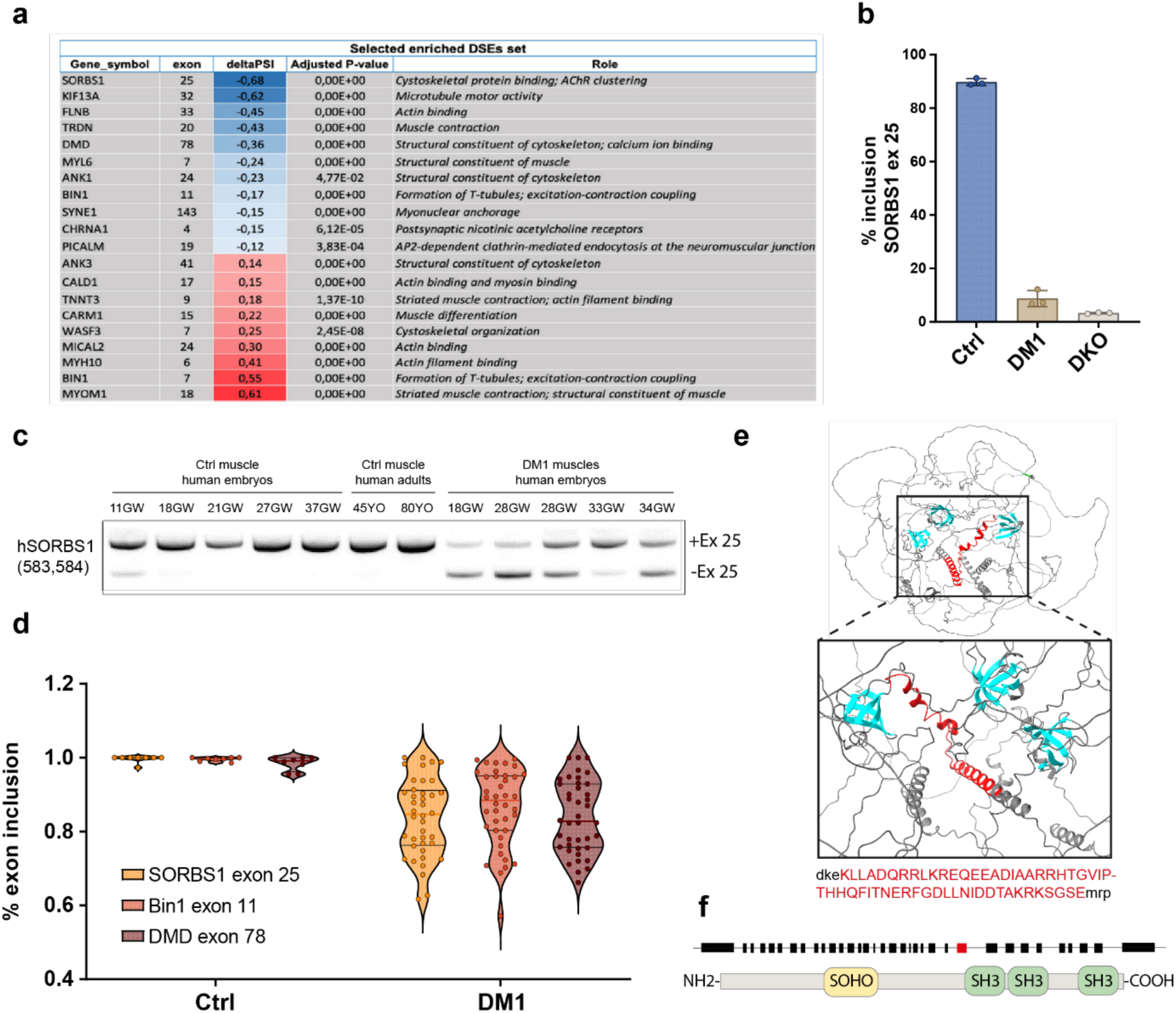
SORBS1 exon 25 is misregulated in skeletal muscle biopsies from DM1 patients. (**a**) List of 30 enriched DSEs identified in common in DM1, DKO MBNL1/2 and WT hiPSC-derived skeletal muscle cells in bulk transcriptomic analysis. For alternative splicing analysis, exons have been numbered according to FasterDB database. (**b**) Bar plots of SORBS1 exon 25 inclusion in hiPSC-derived skeletal muscle cells from WT, DM1 and DKO MBNL1/2. (**c**) RT–PCR analysis of SORBS1 exon 25 alternative splicing in human skeletal muscle samples from control (CTL) and congenital DM1 fetuses (DM1). (**d**) Violin plots of SORBS1 exon 25, BIN1 exon 11, and DMD exon 78 inclusion in tibialis anterior muscles samples from control and DM1 patients obtained from the publicly available DMseq database. Ctrl (N=11); DM1 (N=40). (**e**) AlphaFold analysis of the SORBS1 tridimensional structure. The sequences highlighted in red corresponds to the exon 25. (**f**) Schematic diagram of the SoHo domain and the three SH3 domains of the SORBS1 protein. Black squares represent coding exons, and the red box corresponds to the exon 25.

To assess whether *SORBS1* exon 25 undergoes developmental regulation in human skeletal muscles and whether it is misregulated in DM1 patients, exon 25 inclusion was measured in skeletal muscle samples from control and DM1 fetuses. In control samples, the inclusion of *SORBS1* exon 25 progressively increased during development, with full inclusion after 11 gestational weeks (GW) highlighting its significance during early skeletal muscle formation (**Fig. 1c**). In sharp contrast, *SORBS1* exon 25 remained misregulated throughout skeletal muscle development in congenital DM1 fetuses, failing to reach full inclusion even at 28 and 34 GW (**Fig. 1c**). The Λ1PSI for exon 25 in congenital DM1 fetal skeletal muscles was significantly reduced, with a mean value of 52,6 ± 10% (**Supplementary Fig. 1a**). This finding underscores a persistent developmental defect in splicing regulation in DM1.

To confirm the misregulation of SORBS1 exon 25, the analysis was extended to a larger cohort of DM1 patient samples. Using publicly available RNA sequencing data from the DMseq database (http://www.dmseq.org), and consistent with our initial findings, *SORBS1* exon 25 exhibited significant misregulation across *tibialis anterior* (TA) muscle biopsies from adult DM1 patients with a Λ1PSI of 15,8 ± 3,7% (**Fig. 1d).** This level of misregulation was comparable to that observed for *BIN1* exon 11 (13,1 ± 3,7%) and *DMD* exon 78 (14,5 ± 3,5%), two well-characterized MBNL regulated alternative splices and known to be deregulated in DM1 (**Fig. 1d)**^8,10^.

### Splicing misregulation of *SORBS1* exon 25 impairs mice NMJ stability

To investigate the potential role of SORBS1 exon 25 inclusion, *in silico* modeling was initially performed with AlphaFold2^22^ to predict its localization and possible impact on the protein structure (**Fig. 1e**). The predicted structure showed that exon 25 codes for two εξ-helices, located just upstream of the first SRC homology (SH3) domain, suggesting that this region could be critical for the functional conformation of the SORBS1 protein (**Fig. 1e-f**). To investigate the physiological consequences of *SORBS1* exon 25 mis-splicing in DM1, we employed an exon-skipping strategy. We designed antisense oligonucleotides (ASOs) to target specific sequences in *SORBS1* pre-mRNA and promote exon 25 exclusion (**Supplemental Fig. 2a**). ASOs efficacies were assessed by quantifying exon 25 inclusion 48h post-transfection in human primary skeletal muscle cells (**Supplementary Fig. 2b**). Among the different ASO tested, only one successfully induced an exclusion level comparable to that observed in DM1 skeletal muscle cells, by targeting a putative exonic splicing enhancer (ESE25). To determine whether ASO treatment influenced overall *SORBS1* expression level, we measured total *SORBS1* mRNA levels by quantitative PCR (qPCR) (**Supplementary Fig. 2c-d**). Interestingly, ASO-mediated exon 25 exclusion did not alter total *SORBS1* mRNA expression but selectively reduced the number of transcripts containing exon 25 (**Supplementary Fig. 2c-d**). We then engineered a modified U7 small nuclear RNA (U7-snRNA) construct harboring the antisense sequence targeting ESE25 (*U7-SORBS1-ESE25*) and cloned it into an adeno-associated virus (AAV) (**Fig. 2a**). The left TA muscles of wild type eight-weeks old mice were then injected with *U7-SORBS1-ESE25* virus, whereas the right contralateral muscles were injected with PBS. Muscles were analyzed 2- and 6-months post-injection to evaluate the short- and long-term effects of *SORBS1* exon 25 mis-regulation (**Fig. 2b**).

**Figure 2.**
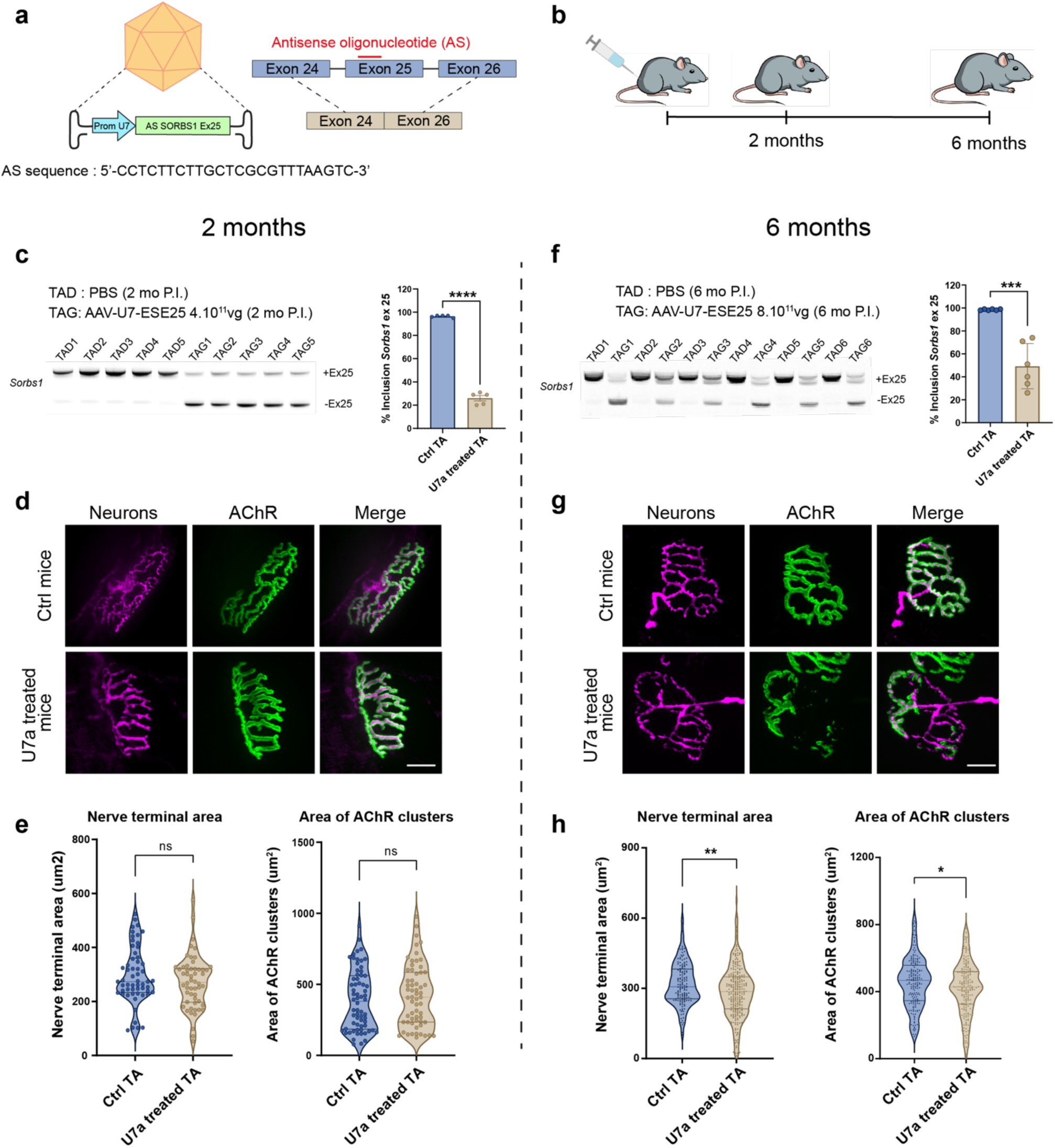
Exclusion of *Sorbs1* exon 25 in mice skeletal muscle affects NMJ stability. (**a**) Schematic representation of the U7 - *Sorbs1* exon 25 antisense constructs (U7-ESE25). (**b**) Schematic representation of the experimental protocol. (**c**) RT-PCR analysis and quantification of *Sorbs1* exon 25 inclusion in TA muscles injected with AAV-U7-ESE25 compared with contralateral TA muscles injected with PBS (Ctrl) 2 months post injection. p < 0,0001, N = 5 mice, Student’s t-test (**d**) Representative immunofluorescence from confocal Z projections of NMJ from TA muscles injected with AAV-U7-ESE25 compared with contralateral TA muscles injected with PBS (Ctrl) 2 months post injection. (**e**) Violin plots of nerve terminal area and AChR area output using NMJ-Morph. Ctrl TA (n=57) from N = 5 mice; U7 treated TA (n=54) from N = 5 mice. (**f**) RT-PCR analysis and quantification of Sorbs1 exon 25 inclusion in TA muscles (N=5) injected with AAV-U7-ESE25 compared with contralateral TA muscles injected with PBS (Ctrl) 6 months post injection. (**g**) Representative immunofluorescence from confocal Z projections of NMJ from TA muscles injected with AAV-U7-ESE25 compared with contralateral TA muscles injected with PBS (Ctrl) 6 months post injection. (**h**) Violin plots of nerve terminal area and AChR area output using NMJ-Morph. Ctrl TA (n=234) from N = 6 mice; U7 treated TA (n=327) from N = 6 mice. p < 0.01 and p <0.05, Student’s t-test.

Expression of *U7-SORBS1-ESE25* in TA mouse muscles reproduced the exon 25 mis-splicing observed in DM1 embryonic muscle biopsies, leading to reduction in exon inclusion of 70,4% ± 2,3% and 49,2 ± 8,1% at 2- and 6-months post-injection, respectively (**Fig. 2c, 2f**). Despite these alterations, no significant defects were observed in the injected TA muscles at 2 months post-injection. Hematoxylin and eosin (H&E) staining of muscle tissue revealed no obvious structural abnormalities in muscle fibers and immunofluorescence analysis of NMJ structure showed no significant changes in NMJ morphology and organization (**Fig. 2d-e and Supplementary Fig. 3a**). Nonetheless, transmission electron microscopy (TEM) analysis uncovered distinct structural abnormalities such as: (i) a reduction in the number of post-junctional folds, indicating compromised post-synaptic membrane integrity, (ii) the invasion of the primary synaptic cleft by terminal Schwann cell processes, and (iii) the presence of degenerative structures within the post-synaptic compartment in TA muscles injected with *U7-SORBS1- ESE25* (**Fig.3a-c**).

By 6 months post-injection, immunofluorescence analysis revealed a fraction of NMJ with pronounced reduction in innervation as evidenced by a significant decrease in the average nerve terminal area (10,5% ± 3,4%; **Fig. 2h**). Further analysis also revealed a significant reduction in AChR receptor area, suggesting progressive deterioration in NMJ integrity (5,7% ± 2,5%; **Fig. 2h**). These observations were corroborated by TEM analysis which demonstrated pronounced structural impairments, including highly disorganized junctional folds with enlarged secondary clefts and numerous degenerative vesicles (**Fig.3e-f**). Additionally, ultrastructural analysis also revealed significant endplate denervation and smaller presynaptic boutons in treated TA muscles (**Fig. 3g-h**). Interestingly, the sarcomere organization of TA muscles remained intact in both 2- and 6- month post injection, with no observed changes in muscle fiber ultrastructure (**Fig. 3d and 3i**). These changes suggest that loss of the 56 amino acids encoded by the exon 25 of *SORBS1* significantly disrupts the stability of the NMJ leading to progressive degeneration over time.

**Figure 3.**
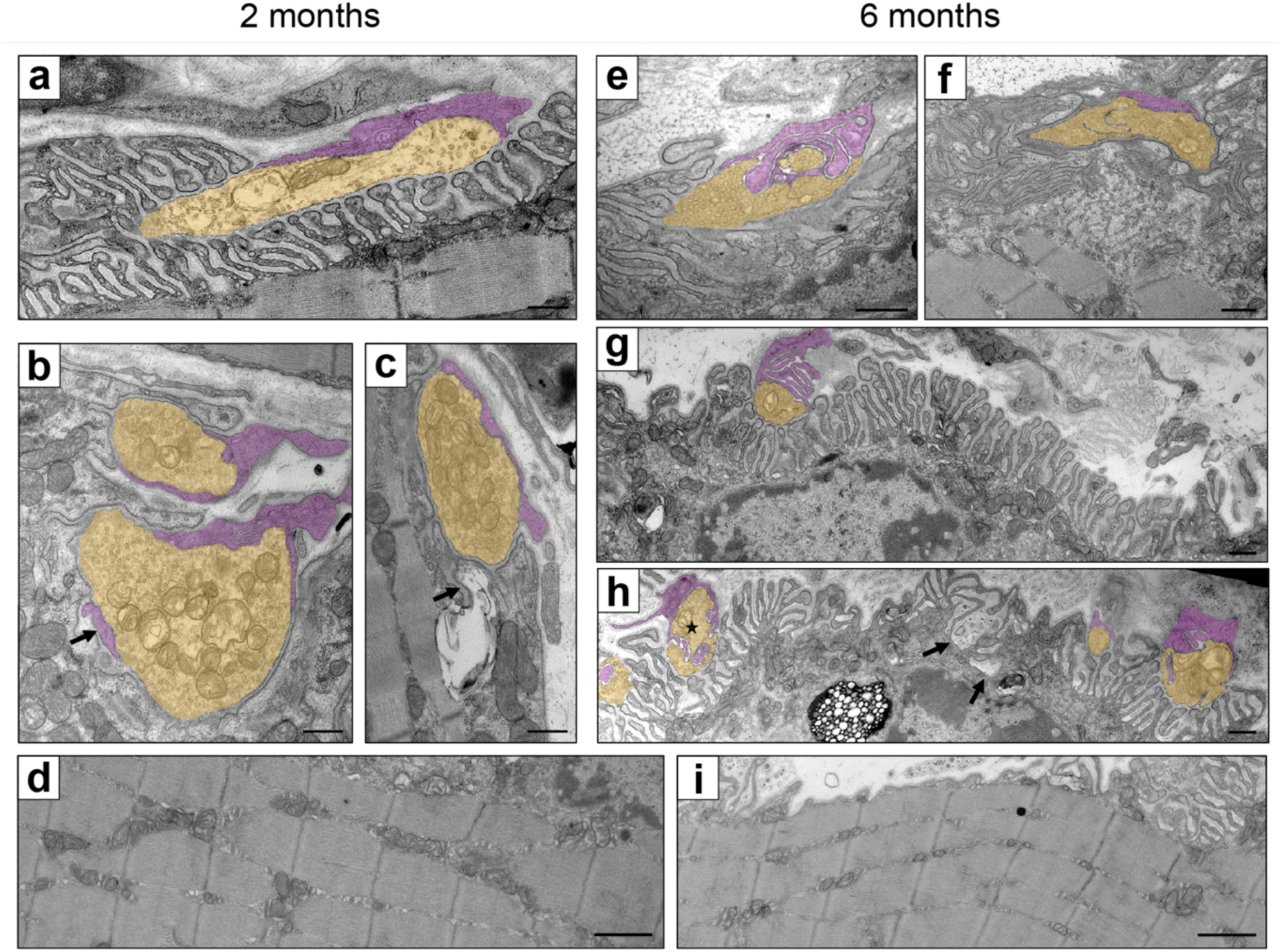
Exclusion of *Sorbs1* exon 25 in mice skeletal muscle disorganize NMJ ultrastructural organization. (**a**) & (**d**) are Ctrl treated tibialis; (**b**), (**c**), (**e**) and (**f**) are U7-ESE25 treated tibialis. (**a**) NMJ from Ctrl treated tibialis at 2 months post injection. The upper part of the neuronal presynaptic bouton is recovered by a terminal Schwann cell process and faces numerous well-ordered parallel post-synaptic junctional folds. (**b-c**) NMJ from U7-ESE25 treated tibialis at 2 months post injection. Presynaptic boutons appear normal with numerous synaptic vesicles and mitochondria, while there is a reduction of post-junctional fold density with rare and short junctional folds. In (**b**) a terminal Schwann cell process intrudes into the primary synaptic cleft (arrow), in (**c**) the arrow points a degenerative structure in the post-synaptic zone. (**d**) Sarcomeres from Ctrl treated TA muscles at 6 months post injection. (**e**-**h**) NMJ from U7-ESE25 treated tibialis at 6 months post injection. Marked endplate denervation with small presynaptic boutons facing only a small portion of the numerous postsynaptic folds. In (**h**) a neuronal bouton is invaded by terminal Schwann cell processes (asterisk). Arrows point to a group of disorganized junctional folds with an abnormal enlarged secondary cleft full of degenerative vesicles. (**i**) Sarcomeres from U7 treated TA muscles at 6 months post injection. Presynaptic terminals are pseudo-colored in yellow and terminal Schwann cell are pseudo-colored in purple. For (**a-c** & **e-h**) scale bars= 500 nm. For (**d** & **i**) Scale bars= 1000 nm

### Exclusion of *sorbs1* exon 25 during zebrafish development impairs locomotor phenotype

To investigate the consequences of *SORBS1* exon 25 exclusion during the neuromuscular development, the exon-skipping strategy was employed in zebrafish embryos (**Fig. 4a**). The *sorbs1* gene in zebrafish consists of 25 exons, with exon 17 corresponding to exon 25 in the human gene. To prevent the developmental splicing switch of *sorbs1*, antisense morpholinos (MO) targeting the splicing site of *sorbs1* exon 17 were injected into one-cell-stage zebrafish embryos. PCR analysis at 50 hours post fertilization (hpf) confirmed efficient exon 17 exclusion, resulting in a ΔPSI reduction of 72.1 ± 5.2% (**Fig. 4b**).

**Figure 4.**
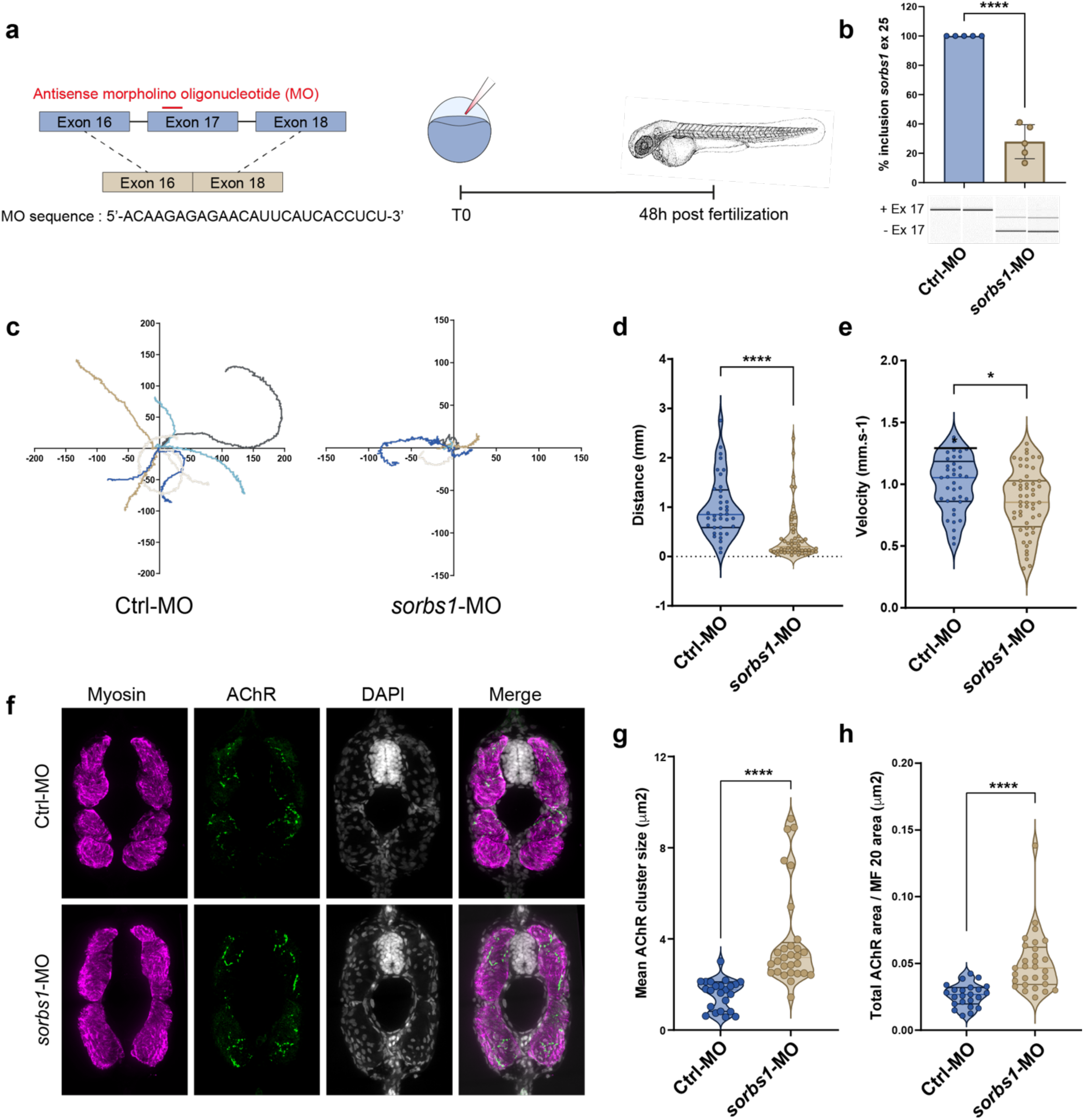
Exclusion of sorbs1 exon 25 during zebrafish development impairs locomotor phenotype. (**a**) Schematic representation of the exon skipping strategy using antisense morpholino oligonucleotide (MO). Zebrafish eggs are microinjected with 0.6 mM of morpholino, and analysis are realized at 48 hours post fertilization (hpf). (**b**) RT-PCR analysis and quantification of sorbs1 exon 25 inclusion on total RNA extracts isolated from whole Ctrl-MO and sorbs1-MO embryos (48hpf); p<0.0001, N = 5, Student’s t-test. (**c**) Color coded video recorded tracks of swim trajectories of individual Ctrl-MO and sorbs1-MO embryos stimulated in touch-evoked escape response (TEER) assay. (**d**) Violin plots representing distribution of swimming distance in TEER assay measured in cm, Ctrl-MO (N=37); sorbs1-MO (N=52) from 3 independent experiments p < 0.001 for distance, p < 0.05 for velocity, Student’s t-test. (**e**) Violin plots representing distribution of swimming velocity in TER assay measured in mm.s^-1^, Ctrl-MO (N=37); sorbs1-MO (N=52) from 3 independent experiments. (**f**) Representative immunofluorescence of cryosectioned 48hpf zebrafish embryos. Images are confocal Z projections from Ctrl-MO and sorbs1-MO zebrafish. (**g**) Violin plots representing distribution of AChR clusters size in μm^2^; Ctrl-MO (n=23) from at least 6 fish; sorbs1-MO (n=29) from at least 6 fishes from 3 independent experiments. p<0.001, Student’s t-test. (**h**) Violin plots representing distribution of AChR area normalized by skeletal muscle area in μm^2^; Ctrl-MO (n=23) from at least 6 fish; sorbs1-MO (n=29) from at least 6 fishes from 3 independent experiments. p<0.001, Student’s t-test.

The swimming trajectory, distance and velocity, were significantly reduced in sorbs1-MO zebrafish compared to controls in the touch-evoked escape response (TEER) test (57.9% ± 12% for distance & 14% ± 0.5% for velocity; **Fig. 4c-e**, **Supplementary Fig. 4a and Supplementary Video 1**). These findings demonstrate that exclusion of *sorbs1* exon 17 during neuromuscular development severely impairs locomotor function.

Immunofluorescence analysis of AChR cluster revealed alterations in NMJ organization (**Fig. 4f**). In *sorbs1*-MO larvae, AChR clusters appeared larger and aggregated in comparison to *mock-* injected larva, where AChR clusters were typically small and dispersed along the muscle fibers (**Fig. 4f**). Quantitative analysis showed an increase in the mean size of individual AChR clusters (147% ± 29,7%; **Fig. 4g**), as well as significant expansion in the total AChR area normalized to skeletal muscle area (92,3% ± 19,0%; **Fig. 4h**). Examination of muscle tissues indicated that although the “U” shaped myosepta remained intact in both control- and *sorbs1*-MO fishes, exon 17 exclusion caused a marked disruption in muscle fibers alignment (**Supplementary Fig. 4b**). This alteration was quantified by a reduction in proper fiber orientation, as measured by local anisotropy (24,32% *±* 5,4%; **Supplementary Fig. 4c**). In addition to its role in maintaining NMJ stability as observed in mice, these findings emphasize that the exclusion of SORBS1 exon 25 during neuromuscular development results in motor impairments and disrupted AChR clustering.

### SORBS1 exon 25 inclusion is mandatory to generate large acetylcholine receptor clusters

Since misregulation of *SORBS1* exon 25 disrupts NMJ formation in both mouse and zebrafish, we hypothesized that similar effects would occur in AChR clustering in human myotubes in response to agrin stimulation. To test this, hiPSC-derived skeletal muscle cells were transfected with ASO sequence targeting the ESE25 region during the final stages of myogenic differentiation (**Fig. 5a**), specifically on day 5 when the myotube formation is initiated.

**Figure 5.**
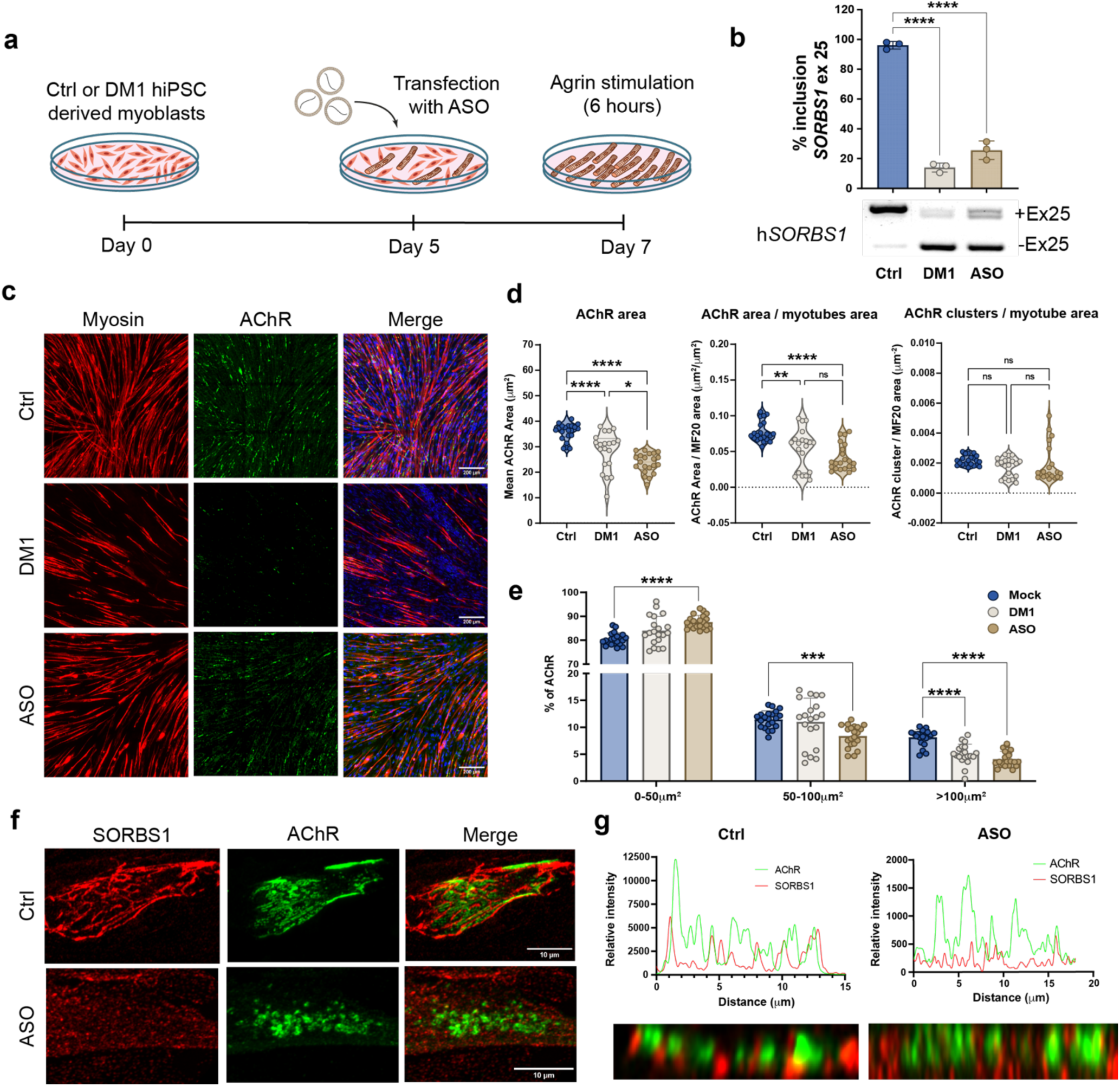
SORBS1 correct splicing is mandatory to generate large AChR clusters. (**a**) Schematic representation of the exon skipping strategy using antisense oligonucleotide (ASO). hiPSCs derived myotubes from controls are terminally differentiated and transfected with 50 nM of ASO at 5 days. At day 7 of differentiation, hiPSC-derived myotubes are treated with 0,5 mg/ml of agrin for 6 hours. (**b**) RT-PCR analysis and quantification of SORBS1 exon 25 inclusion on total RNA extracts isolated from Ctrl, DM1 and ASO transfected cell lysates. (**c**) Representative immunofluorescence of hiPSC derived myotubes at 7 days of differentiation and after 6 hours of agrin stimulation (0,5 mg/ml). Images are confocal Z projections. (**d**) Violin plots representing distribution of AChR clusters area in mm^2^; Ctrl (n=21) from 3 independent experiments; DM1 (n=21) from 3 independent experiments; ASO (n=21) from 3 independent experiments. p < 0.01 and p < 0.0001, one-way-ANOVA. (**e**) Bar plots representing distribution of AChR cluster sizes divided into groups of 0 to 50 μm^2^, 50 to 100 μm^2^ and superior to 100 μm^2^. N=21 from 3 independent experiments. p < 0.001 and p < 0.0001, two-way-ANOVA (**f**) Representative immunofluorescence of AChR clusters in Ctrl and ASO transfected hiPSC derived myotubes in high resolution confocal microscopy. (**g**) Fluorescence intensity profiles of AChR and SORBS1 in newly formed clusters following 6 hours of agrin stimulation.

Transfection with ASO successfully reproduced the exon 25 mis-splicing observed in hiPSC-derived skeletal muscle cells from DM1 patients, leading to a ΔPSI reduction of 82.2% *±* 3.2% and 70.6% *±* 3.5% for DM1 and ASO, respectively (**Fig. 5b**). Immunofluorescence analysis of AChR clusters at 7 days of differentiation, preceded by 6 hours of stimulation with 0.5 μg/ml of agrin, revealed notable alterations in AChR cluster sizes (**Fig. 5c**). Quantitative analysis showed a reduction of 34% *±* 4,5% in ASO-treated hiPSC-derived skeletal muscle cells (**Fig. 5d**). Strikingly, a similar decrease in the mean size of AChR cluster was observed in DM1 hiPSC-derived skeletal muscle cells (**Fig. 5d**). Previous studies have demonstrated that skeletal myoblasts isolated from DM1 patients show reduced fusogenic capacity and diminished ability to form large and mature myotubes^17,23^. To specifically isolate the effect of *SORBS1* exon 25 exclusion on AChR clustering, independent of fusogenic capacity, the total AChR area was normalized to the total myotube area. This analysis revealed reductions of 31,3% *±* 8,4% and 46,9% ± 8,3% in the total AChR area in DM1 and ASO-treated skeletal muscle cells, respectively (**Fig. 5d**). Interestingly, the total number of AChR cluster normalized to the total myotube area remained unchanged in both DM1 and ASO-treated skeletal muscle cells (**Fig. 5d**).

To further characterize the specific population of AChR clusters affected by exon 25 exclusion, the clusters were stratified into three size categories: 0-50 μm^2^, 50-100 μm^2^, and >100 μm^2^ (**Fig. 5e**). A significant reduction was observed in the number of very large AChR clusters (>100 μm^2^) in both DM1 and ASO-treated hiPSC-derived skeletal muscle cells, suggesting that *SORBS1* exon 25 exclusion primarily disrupts AChR cluster size and organization rather than overall cluster formation. Specifically, the total number of clusters >100 μm^2^ was reduced by 38.4% *±* 6.9% and 49.9% *±* 4.2% in DM1 and ASO-treated cells, respectively (**Fig. 5e**). To validate the effects of *SORBS1* exon 25 exclusion on the clustering of AChR in an independent model, we performed experiments in primary control skeletal muscle cells derived from children quadriceps biopsies (**Supplementary Fig. 5a-b**). These analyses confirmed a significant decrease in the total AChR cluster area normalized to myotubes area, with a reduction of 59.8% ± 18.9%, supporting the role of *SORBS1* exon 25 in regulating the clustering of AChR on the myotube membrane.

Using high-resolution imaging with a LSM880 microscope equipped with Airyscan module, we observed distinct spatial interdigitation between SORBS1 and AChR at the plasma membrane in control skeletal muscle cells (**Fig. 5f**). Detailed examination revealed that SORBS1 localization at the plasma membrane was altered upon exon 25 exclusion, transitioning from large, packed protein clusters to more dispersed and fragmented clusters (**Supplementary Fig 5.c**). Notably, in ASO-treated hiPSC derived skeletal muscle cells with mis-spliced exon 25, the typical juxtaposition of SORBS1 and AChR clusters was disrupted, resulting in a shift from “plaque-like” AChR formations to more aggregated, irregular clusters (**Fig. 5f**). Scanning the immunofluorescence intensity across the AChR clusters confirmed this interdigitation pattern in control cells, while ASO-treated cells exhibited significantly reduced fluorescence intensity, indicating a reduction in the presence of protein at the membrane (**Fig. 5g**). These findings indicates that *SORBS1* exon 25 mis-splicing impairs the formation of large, organized AChR clusters in skeletal muscle cells, which may be critical for NMJ stability and proper neuromuscular function in DM1.

## DISCUSSION

This study demonstrates that *SORBS1* exon 25 inclusion is essential for NMJ formation and stability in both animal models and human skeletal muscle cells. Notably, the alternative splicing of *SORBS1* exon 25 is dysregulated in skeletal muscles of DM1 patients, which may contribute to the neuromuscular impairments associated with the disease.

In skeletal muscle biopsies from human control fetuses, *SORBS1* exon 25 inclusion was first observed at 11 gestational weeks (**Fig. 1**), which coincides with the formation of the first NMJs during human embryogenesis^28^. This temporal correlation suggests a potential connection between the developmental regulation of *SORBS1* exon 25 inclusion and the establishment of neuromuscular connectivity. The inclusion of *SORBS1* exon 25 results in the addition of 54 amino acids, forming two εξ-helices upstream of the protein’s first SH3 domain. SORBS1, also known as ponsin or CAP, is an adaptor protein containing three SH3 domains that mediate interactions with various partners. Notably, SORBS1 binds Cbl, playing a critical role in the insulin signaling pathway^29,30^. It localizes to cell-extracellular matrix and cell-cell adhesion sites, where it interacts with proteins such as afadin, vinculin and paxillin^19,20^. Depletion of SORBS1 in HEK293 cells resulted in cytoskeletal disorganization, increasing cell spreading ability associated with an increased cell migration^18^. Beyond these roles, SORBS1 biological functions were also largely associated with cancers, as several SORBS1 mutations have been observed in different studies conducted on human cohorts^31–33^.

In heart and skeletal muscles, SORBS1 has been described to colocalize with vinculin in costameres, where it facilitates the interaction between the cell membrane and the filamin C^34–36^. SORBS1 appears to forge the links between a contractile actin cytoskeleton and a complex protein scaffolding at the cellular membrane. Furthermore, SORBS1 has been identified to interact with the CRK-like protein (CRK-L) in mouse skeletal muscle cells^21^. Its depletion through RNA interference approaches led to the disruption of AChR cluster formation after agrin stimulation^21^. Our findings build on these observations by revealing a new role for SORBS1 in the spatial organization and integrity of the neuromuscular synapse, in addition to its established functions in the structural organization of skeletal muscle.

Exon 25 of *SORBS1* is highly conserved across the different transcripts expressed in skeletal muscle tissues, underscoring a critical function for this coding sequence^37^. Our results demonstrate that increased skipping of SORBS1 exon 25 leads to NMJ destabilization attested by the loss of post-synaptic folds organization and consequently motoneuron denervation. Furthermore, the presence of Terminal Schwann cells within the synaptic clefts of affected NMJ suggests a loss in cellular membrane contact integrity due to the loss of exon 25 inclusion. Forcing *SORBS1* exon 25 exclusion in zebrafish embryos led to strong impaired locomotor phenotypes associated with disorganized AChR clusters. Notably, immunofluorescence analyzes in zebrafish revealed enlarged AChR clusters, contrasting with findings in mouse and human skeletal muscle cell models. The larger AChR clusters observed in zebrafish may reflect compensatory mechanisms similar to those reported following loss of agrin-LRP4 signaling during zebrafish development^38^. The agrin signaling, mediated by the MUSK-LRP4 receptor tyrosine kinase complex, plays a key role in AChR clustering^24,39^. The downstream signaling cascade during the early minutes following agrin stimulation involves the activation of the Rac1 GTPase, PAK1 and actin cytoskeleton reorganization^24–26,40,41^. Other signaling pathways such as GSK3β phosphorylation following AKT signaling has also been described as being implicated in CLASP2 stabilization at microtubule growing ends^27^. However, the protein interactions of SORBS1 and associated signaling, as well as the molecular consequences of its mis-splicing remain to be fully elucidated.

NMJ formation is tightly orchestrated by a cascade of signal exchanges between motoneurons and skeletal muscle fibers. Among the pathways involved, the Agrin/ LRP4/ MuSK/ Dok7 signaling cascade is one of the best-characterized mechanisms critical for NMJ development^42^. Mutations in these genes have been implicated in congenital neuromuscular disorders including congenital myasthenic syndrome (CMS), although most of CMS cases result from mutations in AChR subunits^43^. In addition, some CMS cases arise from mutations affecting pre-mRNA splicing^44^ suggesting that splicing factors and their regulated alternative splicing events may play a critical role in NMJ formation and maintenance. For instance, the splicing factor SRSF1 has been implicated in NMJ development as SRSF1 deficient mice fail to form mature NMJs likely due to impaired regulation of the alternative splicing of the CHRNE and AChR subunits^45^.

Neuromuscular junction dysfunction has been documented in DM1 with neurophysiological studies across different patient cohorts identifying abnormalities in neuromuscular transmission^46,47^. Despite this, the molecular mechanisms underlying these phenotypes remain poorly understood. The sequestration of MBNL protein by toxic (CUG)n-repeats RNA leads to widespread splicing misregulation, potentially disrupting key NMJ-associated splicing events. Notably, recent studies have shown that MBNL depletion in motor neurons impairs NMJ maintenance and function although the precise mechanisms of MBNL action remain unclear^16,48^. Our findings extend the contribution of MBNL proteins to the establishment and maintenance of the NMJ at the post-synaptic level. In alignment with this, the HSA^LR^ mouse model of DM1, which expresses a human skeletal actin transgene with expanded CTG repeats, exhibits *SORBS1* exon 25 mis-splicing along with significant NMJ morphological alterations^49–51^. These abnormalities, including disrupted AChR turnover and synaptic destabilization, highlight the detrimental impact of splice defects on NMJ structure and function in DM1. Our results directly link *SORBS1* exon 25 mis-splicing to ultrastructural NMJ defects, such as impaired AChR clustering, emphasizing the critical role of toxic CUG repeats and MBNL protein loss in disrupting NMJ homeostasis in DM1.

In conclusion, this study highlights the critical role of MBNL-dependent regulation of *SORBS1* exon 25 splicing in the formation of stable AChR clusters and the maintenance of ultrastructural NMJ integrity. By associating the misregulation of *SORBS1* exon 25 splicing to NMJ dysfunction, we provide insights into how splicing defects contribute to the neuromuscular impairments observed in DM1. These findings emphasize the broader impact of splicing dysregulation on NMJ homeostasis and the need to explore other splicing events in DM1. Additionally, this study stresses the importance of developing therapies to restore or preserve NMJ integrity. Finally, our results also point to the significance of understanding the interplay between splicing regulation and cytoskeletal organization at the NMJ, which could reveal additional therapeutic targets for stabilizing the synaptic functions.

## METHODS

### Human biopsies

Fetal skeletal muscle samples were obtained from autopsies, in accordance with the French legislation on ethical rules^10^. Control and congenital DM1 muscle samples were obtained from aborted fetuses showing, respectively, no sign of neuromuscular disease (control) and clinical symptoms of congenital DM1 form with large CTGn>1,000 repeats.

### Bioinformatic analysis

RNA-seq data from GSE86356, Tibialis anterior muscle tissue^52^ were analyzed using Human Genome Reference GRCh37.87, STAR aligner (version 2.7.3a^53^) and the Multivariate Analysis of Transcript Splicing (rMATS version 4.1.2^54^) program, to identify alternatively skipped exons (ASE - JC count) between myotonic dystrophy type 1 (DM1) patients and Control. Spliced exon variants, once detected, are significant when they have an adjusted P-value ≤0.05 and a ΔPSI value ≥10%.

### AAV production and titration

Antisense sequences were cloned in a U7snRNA gene using the BsmI restriction sites of the AAV vector plasmid pSMD2-U7-BsmI^11^ to promote targeted exon skipping. Two pSMD2-U7-SORBS1-ESE25 plasmids were obtained by sub-cloning antisense sequences covering a putative exonic splicing enhancer designed to skip exon 25 (168 nucleotides) of the mouse or human *SORBS1* mRNA (mouse antisense: 5’- CCT-CTT-CTT-GCT-CGC-GCT-TAA-GTC; human antisense: 5’- CCT-CTT-CTT-GCT-CGC-GTT-TAA-GTC). AAV1 or AAV9 viral vectors were prepared by tri-transfection in 293 cells using PEI transfection agent and one of the AAV vector plasmid described above (pSMD2-U7- SORBS1-ESE25), the pXX6 plasmid coding for the viral sequences essential for AAV production and the p0001 or p5E18-VD29 plasmid coding for serotype 1 or 9 capsid respectively. Vector particles were purified on iodixanol gradient and concentrated on Amicon Ultra-15 100K columns (Merck-Millipore, USA). The AAV vectors were titrated as viral genomes (vg) per ml by quantitative real-time PCR using ITR2 (Inverted Terminal Repeats) specific primers at a 60°C annealing temperature (forward 5’-CTC- CAT-CAC-TAG-GGG-TTC-CTT-G and reverse 5’-GTA-GAT-AAG-TAG-CAT-GGC) and the MGB Taqman probe 5’-TAG-TTA-ATG-ATT-AAC-CC.

### Mice experiments

Mice studies conform to the French laws and regulations concerning the use of animals for research were approved by an external Ethical committee (approval n°13333-2018013111391590 v2 delivered by the French Ministry of Higher Education and Scientific Research).

Eight-week old female FVB mice (FVB/NRj, Janvier Cat#SC-FVBN-F, France) were anesthetized by isoflurane for AAV-U7-SORBS1-ESE25 intramuscular (IM) administration of two 45µL injections of vector over a period of 24 h using a 30-G needle in the left tibialis anterior muscle (TA). The right contralateral TA muscle was injected with the same volume of PBS as control. Dose-response studies were carried out with IM injections of vector doses ranging from 0.5 to 8 × 10^11^ vg (1-2 mice per group), and the optimal exon skipping doses were defined at 4 × 10^11^ viral genomes (vg) and 8 × 10^11^ vg for AAV1-U7-SORBS1-ESE25 and AAV9-U7-SORBS1-ESE25 respectively. Mice injected with AAV1- U7-SORBS1-ESE25 were sacrificed 2 months post injection (PI) (n = 10), with collection of TA muscles prepared for either electron microscopy (n = 2), RNA extraction (n = 5), or for neuromuscular junction immunofluorescence (n = 3). Mice injected with AAV9-U7-SORBS1-ESE25 were sacrificed 6 months post injection (PI) (n = 6), with collection of TA muscles prepared for either electron microscopy (n = 2), or neuromuscular junction immunofluorescence (n = 4); a piece of all TA from this later group was also flash frozen in liquid nitrogen for RNA extraction. All mice used in this study were maintained on a 12-hour light/dark cycle and received standard diet and water ad libitum at the animal facility of the Sorbonne University, Paris.

For transmission electron microscopy of neuromuscular junctions, mouse tibialis muscles were fixed with 2% paraformaldehyde, 2% glutaraldehyde in 0.1M phosphate buffer (pH 7.4). Samples were postfixed with 2% OsO4 in 0.1 M phosphate buffer (pH 7.4) for 1 h, then dehydrated in a graded series of acetone including a 1% uranyl acetate staining step in 70% acetone, and finally embedded in epoxy resin (EMBed-812, Electron Microscopy Sciences, USA). Ultra-thin (70 nm) sections were stained with lead citrate, observed with a transmission electron microscope operated at 120 kV (JEOL, Japan), and images were recorded with a Xarosa digital camera (EM-SIS, Germany).

For neuromuscular junction immunofluorescence and morphometry, whole TA muscles were dissected and fixed for 1 h in 4% paraformaldehyde in PBS at room temperature, and then washed and kept at 4°C in PBS. TA muscle fibers were then manually isolated in ice-cold PBS, and incubated overnight with 100 mM glycine in PBS at 4°C. Isolated fibers were then washed three times for 10 min in room temperature PBS, followed by permeabilization and blocking for 4 h (4% BSA, 5% goat serum and 0.5% Triton X-100 in PBS). Fibers were incubated overnight at 4°C in blocking solution with a mouse monoclonal antibody against neurofilament deposited to the DSHB by Jessell, T.M. / Dodd, J. (DSHB Hybridoma Product 2H3) used at 0.5 µg/ml, along with a rabbit polyclonal antibody against synaptophysin (Invitrogen #PA527286) used at à 0.5 µg/ml. After four 1 h washes in PBS, fibers were incubated overnight at 4°C with secondary antibodies Alexa Fluor Plus 555 HCA goat anti-mouse IgG (Invitrogen #A32727) and Alexa Fluor Plus 555 HCA goat anti-rabbit IgG (Invitrogen #A32732), as well as Alexa Fluor 488-conjugated α-bungarotoxin (Invitrogen #B13432) in blocking solution. After four 1 h washes in PBS, fibers were mounted in Vectashield H1000 medium (Vector Laboratories). Imaging was performed on a confocal Nikon Ti2 microscope equipped with a motorized stage and a Yokogawa CSU-W1 spinning disk head coupled with a Prime 95 sCMOS camera (Photometrics) and a 100X/1.43 NA oil-immersion objective at 20°C ambient temperature. Fluorophores were sequentially excited using lasers with wavelengths of 488 and 561 nm respectively. Images were acquired as Z-stack and a semi-automated morphometric analysis was performed using the ImageJ software (https://imagej.nih.gov/ij/) and the NMJ-morph macro^55^ to quantify nerve terminal, endplate and the acetylcholine receptor areas.

### RNA extraction and RT-PCR

Total RNA was isolated using TRIZOL reagent (Life Technologies Cat#15596018, France) from mice TA muscle, human samples, or hiPS cell cultures. Total RNA (1μg) was submitted to reverse transcription using MLV reverse transcriptase and oligo dT_12-18_ (Life Technologies, France). PCR was performed using 25ng of cDNA diluted in platinum Taq polymerase mix containing 1.5 mM MgCl2 (Invitrogen, France) and 0.2µM primers in a final volume of 25 µL. Cycling conditions consisted of a polymerase activation step at 94°C for 5 min, followed by 35 cycles of three steps: 30 s denaturation at 94°C, 30 s annealing at the appropriate temperature and 25 s extension at 72°C. For human *SORBS1*- ex25 PCR, forward primer 5’-CCA-GCT-GAT-TAC-TTG-GAA-TCC-ACG-GAA-G and reverse primer 5’-GTT-CTC-CTT-CAT-ACC-AGT-TCT-GAT-CAA-T were annealed at 58°C. For mouse *SORBS1*-ex25 PCR, forward primer 5’-CCA-GCT-GAT-TAC-TTG-GAG-TCC-ACA-GAA-G and reverse primer 5’-GTT-CAC-CTT-CAT-ACC-AGT-TCT-GGT-CAA-TC were annealed at 60°C. Gel electrophoresis of PCR products was performed in 2.5% agarose and images were acquired on a Geni2 gel imaging system (Ozyme, France). The densitometric analysis of PCR bands was realized using ImageJ Software, and the "Percent Spliced In" (PSI) value corresponding to the fraction of mRNAs that contains exon 25 was calculated as the ratio of the density of the exon inclusion band to the sum of the densities of inclusion and exclusion bands, expressed as a percentage.

### Zebrafish experiments

Adult and larval zebrafish (*Danio rerio*) were maintained at Imagine Institute (Paris) facility and bred according to the National and European Guidelines for Animal Welfare. Experiments were performed on wild type zebrafish larvae from AB strains. Zebrafish were raised in embryo medium: 0,6 g/l aquarium salt (Instant Ocean, Blacksburg, VA) in reverse osmosis water 0,01 mg/l methylene blue. Experimental procedures were approved by the National and Institutional Ethical Committees. Zebrafish were staged in terms of hours post fertilization (hpf) based on morphological criteria and manually dechorionated using fine forceps at 24 hpf. All the experiments were conducted on morphologically normal zebrafish larvae.

A morpholino antisense oligonucleotide (MO; GeneTools, Philomath, USA) was used to specifically knockdown the expression of the *sorbs1* zebrafish orthologue. The MO was designed to bind to a splicing region in exon25 (sorbs1-MO). The sequence is 5’-ACA-AGA-GAG-AAC-AUU-CAU-CAC- CUC-U-3’. Control morpholino (control-MO), not binding anywhere in the zebrafish genome, has the following sequence 5’-CCT-CTT-ACC-TCA-GTT-ACA-ATT-TAT-A-3’. MOs were injected at the final concentration 0.6 mM.

Locomotor behavior of 50 hpf zebrafish larvae were assessed using the Touched-Evoked Escape Response (TEER) test. Briefly, zebrafish were touched on the tail with a tip and the escape response were recorded using a Grasshopper 2 camera (Point Grey Research, Canada) at 30 frames per second. Travelled distance and average velocity were quantified frame per frame for each embryo using the video tracking plugin of FIJI 1.83 software.

### Cell lines

The hiPSC lines used in this study have been previously described^17^. Briefly, DM1 hiPS cell lines with >2000 CTG repeats (62c12, XY, passages p8-20 and 61c7, XY, passages 10-20), control hiPS cells (56c2, passages 10-35) and *MBNL1,2* KOs hiPSCs. CHQ cells are primary myoblasts from biopsy of healthy children quadriceps, a gift from the team of Denis Furling (Institute of Myology, Paris).

Human iPSCs were maintained and passaged in StemMACS iPS-Brew XF medium (Miltenyi Biotec) in vitronectin-coated culture dishes (Gibco) as previously described^17^. Skeletal muscle differentiation experiments were performed using the commercially available STEMdiff Myogenic Progenitor Supplement Kit (StemCell Technologies). Briefly, hiPSCs were dissociated using StemPro Accutase Cell Dissociation Reagent (Gibco) and plated at 30,000 cells per cm2 on growth factor-reduced Matrigel (Corning), and the complete medium was changed and replaced with STEMdiff Myogenic Progenitor Medium A on day 0 and day 1. On day 2 and day 3, the complete medium was changed and replaced by STEMdiff Myogenic Progenitor Medium B. On day 4 and day 5, the complete medium was changed and replaced by STEMdiff Myogenic Progenitor Medium C. From day 6 to day 29, the complete medium was daily changed and replaced by STEMdiff Myogenic Progenitor Medium D. On day 30, cells were dissociated with trypsin-EDTA 0.05% (Gibco), amplified for 2 passages, dissociated, banked and cryopreserved. For terminal differentiation, cells were thawed at 40,000 cells per cm2 in MyoCult- SF Expansion Supplement Kit (StemCell Technologies) for 2-3 days. When the cells reached confluence, the medium was replaced with MyoCult Differentiation Kit (StemCell Technologies) and the cells were maintained in culture for 7 days to allow myotube formation. RNA-Seq: Data are available on GEO database (GSE161897).

Human primary CHQ myoblasts were thawed and plated at 40.000 cells per cm^2^ cultivated in a growth medium consisting of DMEM-F12 glutamax (Life Technologies) supplemented with 20% FBS (Sigma-Aldrich), 1% Insulin human solution (Sigma-Aldrich^®^) and 0,1% penicillin/steptomycine (Life Technologies^®^). After 2-3 days, once the cells reached the confluence, myogenic differentiation was induced by switching the cell cultures to DMEM supplemented with 5% horse serum (Life Technologies) for 3 days to allow myotubes formation.

### ASO(s) treatments

Differentiated hiPSCs-derived control myotubes or human primary control myoblasts were transfected at day 5 of differentiation with 50 nM 2-OMe ASOs using RNAi Max transfection reagent (Life Technologies) according to the manufacturer’s protocol. At 48h post-transfection, cells were stimulated with 0.5 µg/ml rat agrin (R&D Systems) for various time periods depending on the experiment, and harvested for analysis.

### RT-PCR and Agilent DNA chips analysis for Alternative Splicing analysis

Total cellular RNA was extracted using the RNeasy Micro/Mini Kit (Qiagen) and reverse transcribed using random hexamers, oligoDT and Superscript III Reverse Transcriptase Kit (Invitrogen) according to the manufacturer’s protocol. For splicing analysis, PCR amplification was performed using recombinant Q5 DNA polymerase (New England Biolabs) and primers listed in Table S1. Amplification was performed with a first step at 95°C for 3’, followed by 35 cycles of 15’’ at 95°C, 30’’ at 60°C, 30’’ at 72°C, with a final extension of 10’’ at 72°C. PCR products were analyzed on agarose gels or Agilent DNA chips and quantified on a BioAnalyzer 2100.

### Gene expression analysis by quantitative PCR

Total cellular RNA was extracted using the RNeasy Micro/Mini Kit (Qiagen) and reverse transcribed using random hexamers, oligoDT and Superscript III Reverse Transcriptase Kit (Invitrogen) according to the manufacturer’s protocol. Quantitative PCR reactions were performed in 384-well plates on a QuantStudio 12K Flex Real-Time PCR System (Applied Biosystems) using Power SYBR Green 2× Master Mix (Life Technologies). Each well contained 2.5 μL of cDNA in a final volume of 10 μL containing the SYBR Green mix and the appropriate primers listed in Table S1. The 2-ΔΔCt method was used to determine the relative expression level of each gene.

### Immunocytochemistry

Cells were fixed for 10’’ in PBS containing 4% PFA and 0.1% Triton X100. Cells were washed three times with PBS and blocked for 1 hour in PBS containing 3% BSA and 0.1% Tween20. Primary antibodies listed in Table S2 were incubated 30” at RT in PBS containing 3% BSA and 0.1% Tween20. Cells were washed three times in PBS-tween20 0.1% and incubated for 30” at RT with appropriate fluorescently labelled secondary antibodies listed in Table S2. Images were acquired using two different microscopes. Zebrafish sections and hiPSC-derived myotubes for AChR cluster quantification were imaged using a confocal spinning disk microscope (Zeiss Axio Observer Z.1) equipped with a CSU-X1 Yokogawa spinning scanning unit and an Orca Fusion-BT camera (Hamamatsu). Images were acquired using a Zeiss Plan-Apo 63x/1.40 NA oil objective or a Zeiss Plan-Apo 20x/0.8 NA air objective. Each wavelength was acquired separately with a Z-step width of 300 nm. Metamorph software was used for image acquisition. High magnification images of AChR clusters were obtained on a scanning confocal microscope (Zeiss LSM 880) using a Zeiss 63X/1.4 Plan-Apo Oil DIC and Airyscan detector mode. Zen black software was used for image acquisition.

### Protein extraction and Western blot analysis

Western blots analyses were performed as previously described (Mérien, 2020). Briefly, cells were lyzed in RIPA 1X buffer (Sigma) containing protease inhibitors (Sigma) and phosphatases inhibitors (Roche). Proteins were quantified by Pierce BCA Protein Assay kit (Pierce). Protein extracts (15 to 20 μg) were loaded on a 4-12% SDS-PAGE gradient (NuPage Bis–Tris gels, Invitrogen) and transferred onto Gel Transfer Stacks Nitrocellulose membranes (Invitrogen) using the iBlot2 Dry Blotting System (Invitrogen). Membranes were then incubated overnight at 4°C with the following primary antibodies: SORBS1 (Abcam, 1:1000). After hybridization of the peroxidase-conjugated secondary antibody (1:10000), immunoreactive bands were revealed by using Amersham ECL Select Western Blotting Detection Reagents (GE Healthcare). Equal protein loading was verified by the detection of β-Actin using the A3854 Monoclonal Anti-β-Actin−Peroxidase antibody (Sigma, 1:10000).

### Statistical analysis

All data were processed using Prism 9®. For statistical analysis, either Student’s t-test or one-way analyses of variance were used as appropriate. Values are represented as mean ± SD. Values of p < 0,05 were considered significant (*p < 0,05; **p < 0,005; ***p < 0,0005; ****p < 0,0001).

## Acknowledgments

We thank the MYOVECTOR technical platform of the Centre of Research in Myology-UMRS974 (Paris, France) for AAV production, and the Penn Vector Core, Gene Therapy Program (University of Pennsylvania, Philadelphia, USA) for providing the pAAV1 (p0001) and pAAV9 plasmids (p5E18- VD29). We also gratefully acknowledge support from the PSMN (Pôle Scientifique de Modélisation Numérique) of the ENS de Lyon for the computing resources.

---

**Supplementary figure 1:**
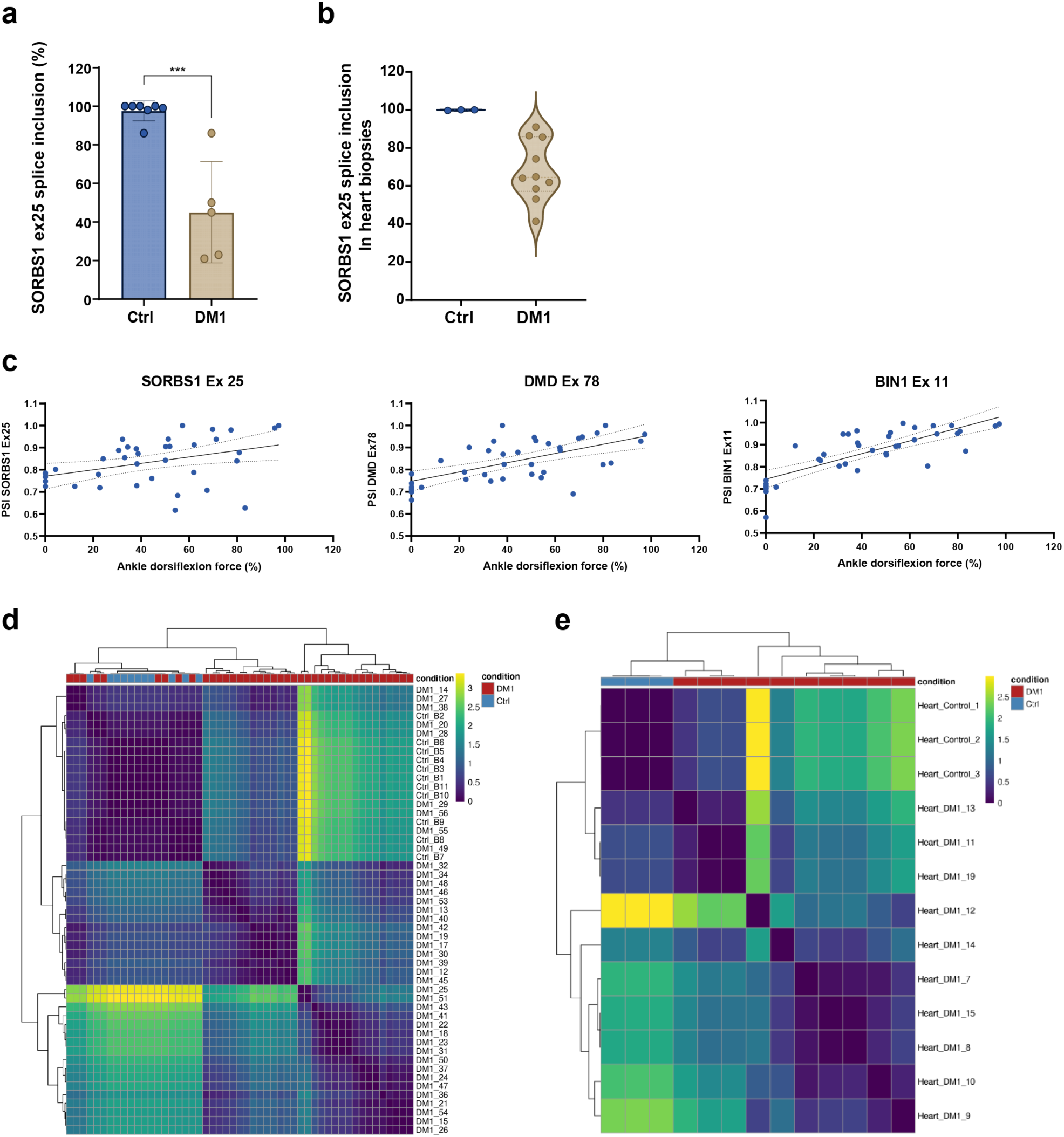
**(a)** Bar graph representing quantification of SORBS1 exon 25 inclusion in skeletal muscle biopsies from human fetuses. p < 0.001, Student’s t-test (**b**) Violin plots of SORBS1 exon 25 inclusion in the heart muscles samples from control and DM1 patients obtained from the publicly available DMseq database. (**c**) Correlation between the splice inclusion of *SORBS1* exon 25, *BIN1* exon 11, and *DMD* exon 78 inclusion in tibialis anterior muscles samples and the ankle dorsiflexion force (%) from the publicly available DMseq database. (**d**) Heatmap indicating the group hierarchy of SORBS1 exon 25 inclusion from the tibialis anterior skeletal muscle samples according to their proximity. (**e**) Heatmap indicating the group hierarchy of SORBS1 exon 25 inclusion from the heart muscle samples according to their proximity.

**Supplementary figure 2:**
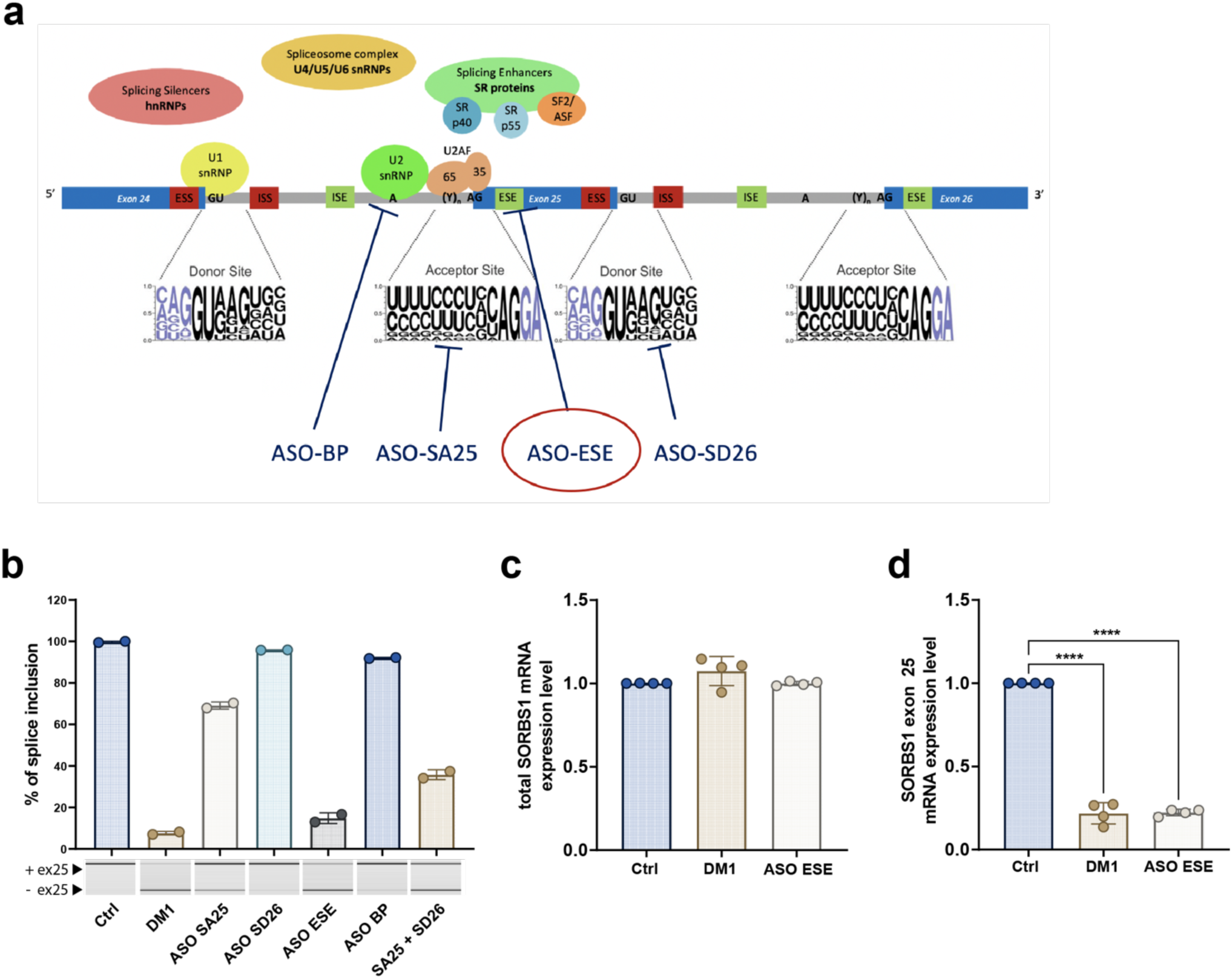
(**a**) Schematic representation of the different antisense oligonucleotide (ASO) tested and their targeted region on *SORBS1* pre-mRNA. ISE: Intron Splice Enhancer, ISS: Intron Splice Suppressor, ESE: Exon Splice Enhancer, ESS: Exon Splice Suppressor, BP: Branching Point, SA: Acceptor Site, SD: Donor Site (**b**) PCR experiments of SORBS1 exon 25 splicing profile in human primary myotubes 48h post transfection (N=2 independent experiments). (**c**) RT-qPCR experiments of total SORBS1 mRNA in Ctrl, DM1 and Ctrl-transfected hiPSC derived myotubes with the ASO-ESE relative to Ctrl. (N = 4 independent experiments). (**d**) RT-qPCR experiments of SORBS1 mRNA containing the exon 25 in Ctrl, DM1 and Ctrl-transfected hiPSC derived myotubes with the ASO-ESE relative to Ctrl. (N = 4 independent experiments, p < 0.0001, one-way ANOVA).

**Supplementary figure 3:**
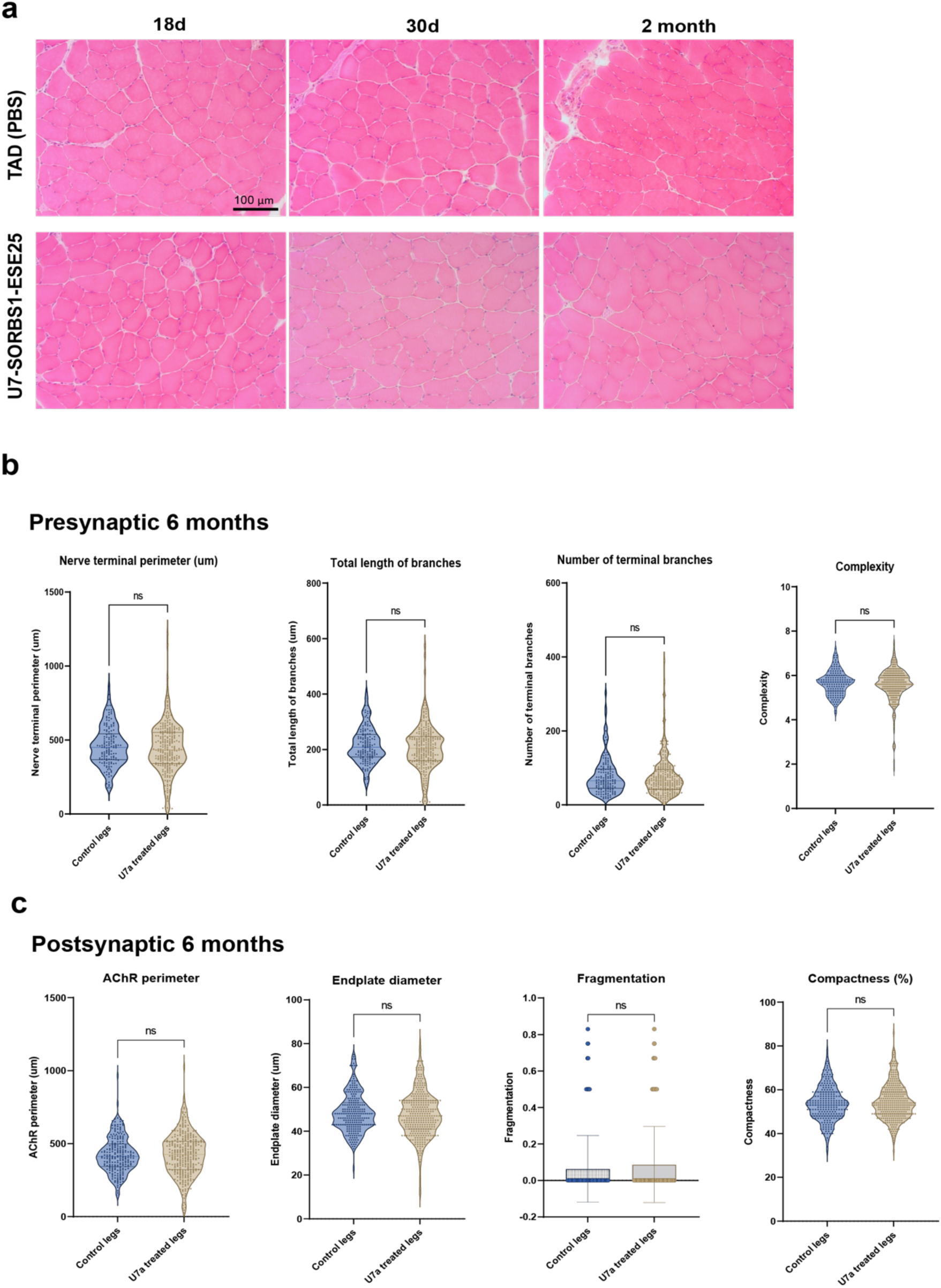
**(a)** Hematoxylin and eosin staining in TA muscles injected either with PBS or AAV-U7-SORBS1-ESE25. (**b**) Violin plots of presynaptic outputs using NMJ-Morph at 6 months post-injection. Ctrl TA (n=234) from N = 6 mice; U7 treated TA (n=327) from N = 6 mice. (**c**) Violin plots of postsynaptic outputs using NMJ-Morph (N=5, n=234) at 6 months post-injection. Ctrl TA (n=234) from N = 6 mice; U7 treated TA (n=327) from N = 6 mice.

**Supplementary figure 4:**
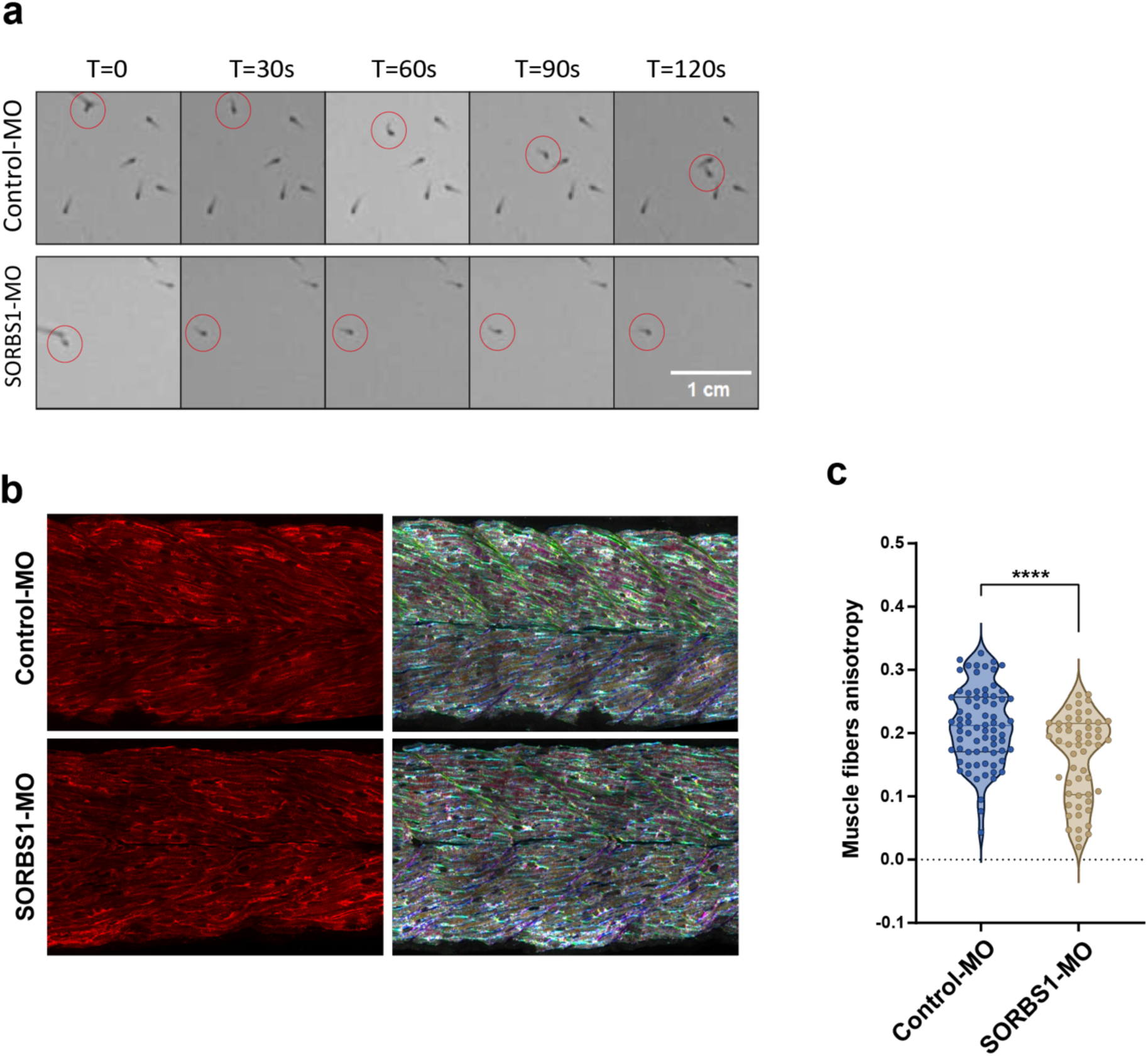
**(a)** Time lapse from recorded videos of swimming comportment of individual Ctrl-MO and *sorbs1*-MO embryos stimulated in touch-evoked escape response (TEER) assay. (**b**) Representative immunofluorescence of cryosectioned 48hpf zebrafish embryos. Images are confocal Z projections from Ctrl-MO and *sorbs1*-MO zebrafish. Muscle fibers orientations are color coded using Orientation J plugin in ImageJ. (c) Violin plots representing distribution of muscle fibers anisotropy; Ctrl-MO (n=23) from at least 6 fish; *sorbs1*-MO (n=29) from at least 6 fish. P < 0.0001, Student’s t-test.

**Supplementary figure 5:**
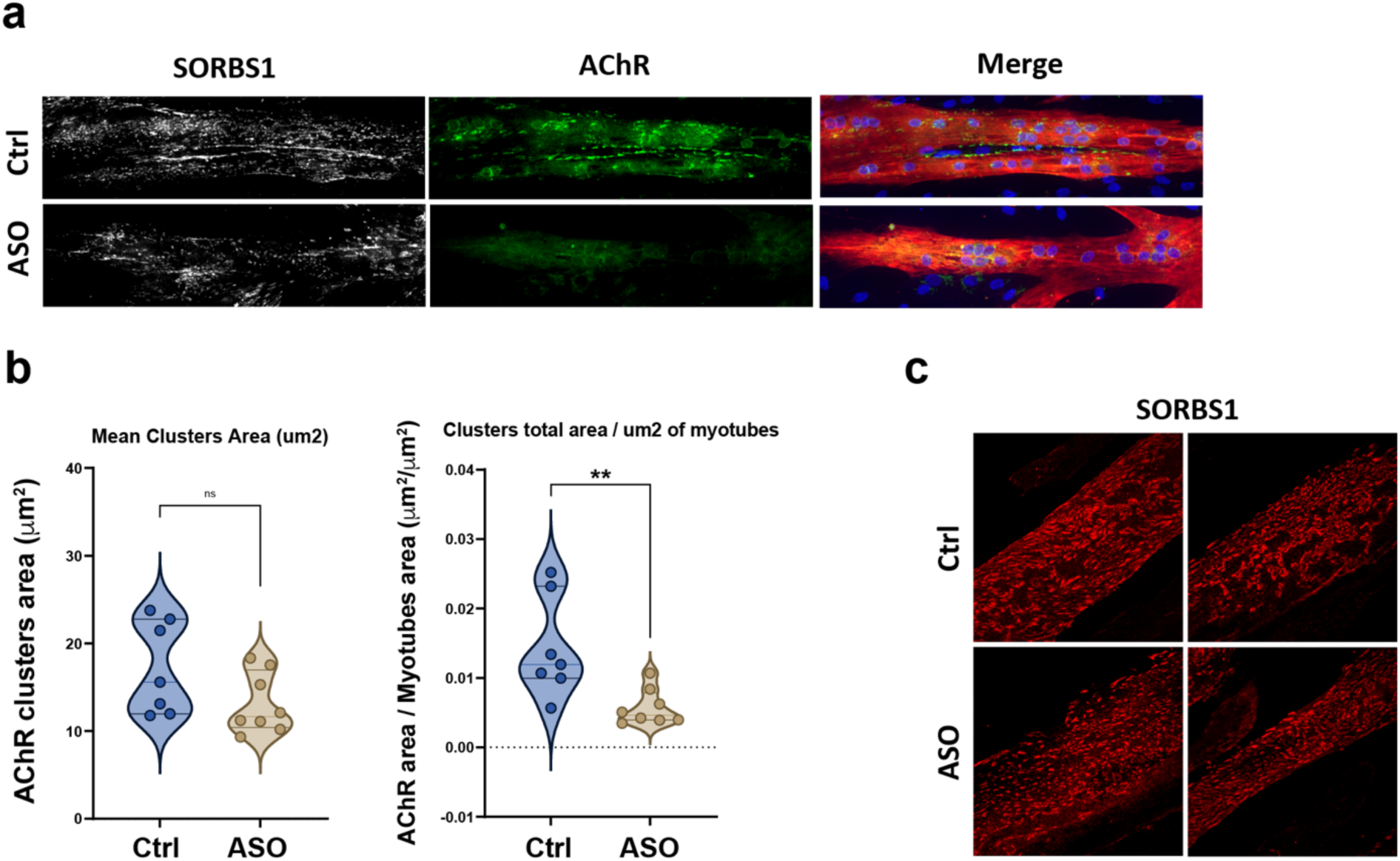
**(a)** Representative immunofluorescence of primary human myotubes at 3 days of differentiation and after 6 hours of agrin stimulation (0,5 mg/ml). Images are confocal Z projections. (**b**) Violin plots representing distribution of AChR clusters area in mm^2^; Ctrl (n=7) from 2 independent experiments; ASO (n=7) from 2 independent experiments. p < 0.01, Student’s t-test. **(c)** Representative immunofluorescence of SORBS1 location on the membrane of primary human myotubes at 3 days of differentiation.

## REFERENCES

1. Yum, K., Wang, E. T. & Kalsotra, A. Myotonic dystrophy: disease repeat range, penetrance, age of onset, and relationship between repeat size and phenotypes. Curr. Opin. Genet. Dev. 44, 30–37 (2017).

2. De Antonio, M. et al. Unravelling the myotonic dystrophy type 1 clinical spectrum: A systematic registry-based study with implications for disease classification. Rev. Neurol. (Paris) 172, 572–580 (2016).

3. Michel, L., Huguet-Lachon, A. & Gourdon, G. Sense and Antisense DMPK RNA Foci Accumulate in DM1 Tissues during Development. PLOS ONE 10, e0137620 (2015).

4. Wheeler, T. M., Krym, M. C. & Thornton, C. A. Ribonuclear foci at the neuromuscular junction in myotonic dystrophy type 1. Neuromuscul. Disord. 17, 242–247 (2007).

5. Thomas, J. D. et al. Disrupted prenatal RNA processing and myogenesis in congenital myotonic dystrophy. Genes Dev. 31, 1122–1133 (2017).

6. Charlet-B., N., et al. Loss of the Muscle-Specific Chloride Channel in Type 1 Myotonic Dystrophy Due to Misregulated Alternative Splicing. Mol. Cell 10, 45–53 (2002).

7. Savkur, R. S., Philips, A. V. & Cooper, T. A. Aberrant regulation of insulin receptor alternative splicing is associated with insulin resistance in myotonic dystrophy. Nat. Genet. 29, 40–47 (2001).

8. Fugier, C. et al. Misregulated alternative splicing of BIN1 is associated with T tubule alterations and muscle weakness in myotonic dystrophy. Nat. Med. 17, 720–725 (2011).

9. Tang, Z. Z. et al. Muscle weakness in myotonic dystrophy associated with misregulated splicing and altered gating of CaV1.1 calcium channel. Hum. Mol. Genet. 21, 1312–1324 (2012).

10. Rau, F. et al. Abnormal splicing switch of DMD’s penultimate exon compromises muscle fibre maintenance in myotonic dystrophy. Nat. Commun. 6, 7205 (2015).

11. Moulay, G. et al. Alternative splicing of clathrin heavy chain contributes to the switch from coated pits to plaques. J. Cell Biol. 219, e201912061 (2020).

12. Nakamori, M. et al. Splicing biomarkers of disease severity in myotonic dystrophy. Ann. Neurol. 74, 862–872 (2013).

13. Jauvin, D. et al. Targeting DMPK with Antisense Oligonucleotide Improves Muscle Strength in Myotonic Dystrophy Type 1 Mice. Mol. Ther. - Nucleic Acids 7, 465–474 (2017).

14. Klinck, R. et al. RBFOX1 Cooperates with MBNL1 to Control Splicing in Muscle, Including Events Altered in Myotonic Dystrophy Type 1. PLoS ONE **9**, e107324 (2014).

15. Lee, K.-Y. et al. Mice lacking MBNL1 and MBNL2 exhibit sudden cardiac death and molecular signatures recapitulating myotonic dystrophy. Hum. Mol. Genet. 31, 3144– 3160 (2022).

16. Frison-Roche, C. et al. MBNL deficiency in motor neurons disrupts neuromuscular junction maintenance and gait coordination. Brain awae336 (2024) doi:10.1093/brain/awae336.

17. Mérien, A. et al. CRISPR gene editing in pluripotent stem cells reveals the function of MBNL proteins during human *in vitro* myogenesis. Hum. Mol. Genet. 31, 41–56 (2021).

18. Zhang, M. et al. CAP interacts with cytoskeletal proteins and regulates adhesion-mediated ERK activation and motility. EMBO J. 25, 5284–5293 (2006).

19. Ichikawa, T. et al. Vinexin family (SORBS) proteins play different roles in stiffness-sensing and contractile force generation. J. Cell Sci. jcs.200691 (2017) doi:10.1242/jcs.200691.

20. Kuroda, M., Ueda, K. & Kioka, N. Vinexin family (SORBS) proteins regulate mechanotransduction in mesenchymal stem cells. Sci. Rep. 8, 11581 (2018).

21. Hallock, P. T., Chin, S., Blais, S., Neubert, T. A. & Glass, D. J. Sorbs1 and -2 Interact with CrkL and Are Required for Acetylcholine Receptor Cluster Formation. Mol. Cell. Biol. 36, 262–270 (2016).

22. Jumper, J. et al. Highly accurate protein structure prediction with AlphaFold. Nature 596, 583–589 (2021).

23. Ludovic, A. et al. Immortalized human myotonic dystrophy muscle cell lines to assess therapeutic compounds. Dis. Model. Mech. dmm.027367 (2017) doi:10.1242/dmm.027367.

24. Luo, Z. G. et al. Regulation of AChR Clustering by Dishevelled Interacting with MuSK and PAK1. Neuron 35, 489–505 (2002).

25. Bai, Y. et al. Balanced Rac1 activity controls formation and maintenance of neuromuscular acetylcholine receptor clusters. J. Cell Sci. jcs.215251 (2018) doi:10.1242/jcs.215251.

26. Klockner, I., Schutt, C., Gerhardt, T., Boettger, T. & Braun, T. Control of CRK-RAC1 activity by the miR-1/206/133 miRNA family is essential for neuromuscular junction function. Nat. Commun. 13, 3180 (2022).

27. Schmidt, N. et al. Agrin regulates CLASP2-mediated capture of microtubules at the neuromuscular junction synaptic membrane. J. Cell Biol. 198, 421–437 (2012).

28. Nebol, A. & Gouti, M. A new era in neuromuscular junction research: current advances in self-organized and assembled in vitro models. Curr. Opin. Genet. Dev. 87, 102229 (2024).

29. Chang, T.-J. et al. Genetic variation of SORBS1 gene is associated with glucose homeostasis and age at onset of diabetes: A SAPPHIRe Cohort Study. Sci. Rep. 8, 10574 (2018).

30. Gong, S. et al. A variation in SORBS1 is associated with type 2 diabetes and high-density lipoprotein cholesterol in Chinese population. Diabetes Metab. Res. Rev. 38, e3524 (2022).

31. Lu, Z. & Gao, Y. Screening differentially expressed genes between endometriosis and ovarian cancer to find new biomarkers for endometriosis. Ann. Med. 53, 1377–1389 (2021).

32. Wang, N. et al. Development and Validation of a Prognostic Classifier Based on Lipid Metabolism-Related Genes for Breast Cancer. J. Inflamm. Res. **Volume** 15, 3477–3499 (2022).

33. Shen, H., Xu, X., Fu, Z., Xu, C. & Wang, Y. The interactions of CAP and LYN with the insulin signaling transducer CBL play an important role in polycystic ovary syndrome. Metabolism 131, 155164 (2022).

34. Gehmlich, K. et al. Paxillin and Ponsin Interact in Nascent Costameres of Muscle Cells. J. Mol. Biol. 369, 665–682 (2007).

35. Gehmlich, K. et al. Ponsin interacts with Nck adapter proteins: implications for a role in cytoskeletal remodelling during differentiation of skeletal muscle cells. Eur. J. Cell Biol. 89, 351–364 (2010).

36. Mandai, K. et al. Ponsin/SH3P12: An l-Afadin– and Vinculin-binding Protein Localized at Cell–Cell and Cell–Matrix Adherens Junctions. J. Cell Biol. 144, 1001–1018 (1999).

37. Lin, W.-H. et al. Cloning, Mapping, and Characterization of the Human Sorbin and SH3 Domain Containing 1 (SORBS1) Gene: A Protein Associated with c-Abl during Insulin Signaling in the Hepatoma Cell Line Hep3B. Genomics 74, 12–20 (2001).

38. Walker, L. J., Roque, R. A., Navarro, M. F. & Granato, M. Agrin/Lrp4 signal constrains MuSK-dependent neuromuscular synapse development in appendicular muscle. Development 148, dev199790 (2021).

39. Kim, N. et al. Lrp4 Is a Receptor for Agrin and Forms a Complex with MuSK. Cell 135, 334–342 (2008).

40. Lee, C. W. et al. Regulation of acetylcholine receptor clustering by ADF/cofilin-directed vesicular trafficking. Nat. Neurosci. 12, 848–856 (2009).

41. Kwan, H.-L. R. et al. Nerve-independent formation of membrane infoldings at topologically complex postsynaptic apparatus by caveolin-3. Sci. Adv. 9, eadg0183 (2023).

42. Ohno, K. et al. Splicing regulation and dysregulation of cholinergic genes expressed at the neuromuscular junction. J. Neurochem. 142, 64–72 (2017).

43. Deenen, J. C. W., Horlings, C. G. C., Verschuuren, J. J. G. M., Verbeek, A. L. M. & Van Engelen, B. G. M. The Epidemiology of Neuromuscular Disorders: A Comprehensive Overview of the Literature. J. Neuromuscul. Dis. 2, 73–85 (2015).

44. Ohno, K. A frameshifting mutation in CHRNE unmasks skipping of the preceding exon. Hum. Mol. Genet. 12, 3055–3066 (2003).

45. Liu, Y. et al. Splicing Factor SRSF1 Is Essential for Satellite Cell Proliferation and Postnatal Maturation of Neuromuscular Junctions in Mice. Stem Cell Rep. 15, 941–954 (2020).

46. Bombelli, F. et al. Neuromuscular transmission abnormalities in myotonic dystrophy type 1: A neurophysiological study. Clin. Neurol. Neurosurg. 150, 84–88 (2016).

47. Krishnan, A. V. & Kiernan, M. C. Axonal function and activity-dependent excitability changes in myotonic dystrophy. Muscle Nerve 33, 627–636 (2006).

48. Tahraoui-Bories, J. et al. MBNL-dependent impaired development within the neuromuscular system in myotonic dystrophy type 1.

49. Lee, K. et al. Compound loss of muscleblind-like function in myotonic dystrophy. EMBO Mol. Med. 5, 1887–1900 (2013).

50. Tomàs, J. et al. Presynaptic Membrane Receptors Modulate ACh Release, Axonal Competition and Synapse Elimination during Neuromuscular Junction Development. Front. Mol. Neurosci. 10, 132 (2017).

51. Falcetta, D. et al. CaMKIIβ deregulation contributes to neuromuscular junction destabilization in Myotonic Dystrophy type I. Skelet. Muscle 14, 11 (2024).

52. Wang, E. T. et al. Transcriptome alterations in myotonic dystrophy skeletal muscle and heart. Hum. Mol. Genet. 28, 1312–1321 (2019).

53. Dobin, A. et al. STAR: ultrafast universal RNA-seq aligner. Bioinformatics 29, 15–21 (2013).

54. Wang, Y. et al. rMATS-turbo: an efficient and flexible computational tool for alternative splicing analysis of large-scale RNA-seq data. Nat. Protoc. 19, 1083–1104 (2024).

55. Jones, R. A. et al. NMJ-morph reveals principal components of synaptic morphology influencing structure–function relationships at the neuromuscular junction. Open Biol. 6, 160240 (2016).

